# Brain-wide neuronal activation and functional connectivity are modulated by prior exposure to repetitive learning episodes

**DOI:** 10.1101/2021.03.28.437394

**Authors:** Dylan J. Terstege, Isabella M. Durante, Jonathan R. Epp

**Affiliations:** Department of Cell Biology and Anatomy, Hotchkiss Brain Institute, Cumming School of Medicine, University of Calgary, 3330 Hospital Drive NW, Calgary, Alberta, Canada T2N 4N1

**Keywords:** Cognitive Stimulation, Functional Connectivity, Spatial learning, Context memory

## Abstract

Memory storage and retrieval are shaped by past experiences. Prior learning and memory episodes have numerous impacts on brain structure from micro to macroscale. Previous experience with specific forms of learning increases the efficiency of future learning. It is less clear whether such practice effects on one type of memory might also have transferable effects to other forms of memory. Different forms of learning and memory rely on different brain-wide networks but there are many points of overlap in these networks. Enhanced structural or functional connectivity caused by one type of learning may be transferable to another type of learning due to overlap in underlying memory networks. Here, we investigated the impact of prior chronic spatial training on the task-specific functional connectivity related to subsequent contextual fear memory recall in mice. Our results show that mice exposed to prior spatial training exhibited decreased brain-wide activation compared to control mice during the retrieval of a context fear memory. With respect to functional connectivity, we observed changes in several network measures notably an increase in global efficiency. Interestingly, we also observed an increase in network resilience based on simulated targeted node deletion. Overall, this study suggests that chronic learning has transferable effects on the functional connectivity networks of other types of learning and memory. The generalized enhancements in network efficiency and resilience suggest that learning itself may protect brain networks against deterioration.

## Introduction

It has been well established that prior learning experiences alter the canvas against which new learning occurs. Learning results in numerous structural changes in the brain ranging from cellular and synaptic changes (Lendvai et al., 2000; Nyberg et al., 2003; Holtmaat et al., 2005; De Paola et al., 2006; Epp et al., 2013) to altered macroscale measurements of regional size and shape (Maguire et al., 2003; Draganski et al., 2004; Bermudez et al., 2008; Hyde et al., 2009; Scholz et al., 2009). A classic example of learning-induced structural changes is the change in hippocampal volume that occurs as a result of intense practice with spatial navigation in London taxi drivers (Maguire et al., 2000). Similar increases in hippocampal volume have also been observed in mice that were trained on a spatial learning task (Lerch et al., 2011).

To the extent that there are relationships between brain function and underlying structure, it should be predicted that learning-induced structural changes should also induce functional changes. Training-induced increases in hippocampal volume for example are also associated with enhanced memory performance (Bohbot et al., 2007).

In addition to structural changes, learning has been shown in some studies to change the organization of memories in the brain. In rats, previous studies have indicated that prior training with a memory task can prevent lesion-induced deficits in both similar and slightly distinct memory tasks (Clark and Delay, 1991; Ocampo et al., 2018). This suggested that prior learning experiences fundamentally change how and where future memories are encoded (Owen et al., 2010; Nouchi et al., 2012; West et al., 2017). Experiments such as these suggest some form of reorganization but do not give a complete picture as to how this reorganization has occurred.

Functional imaging experiments in humans have provided evidence that cognitive stimulation, or memory training, alter brain functional connectivity (Martínez et al., 2013; Dresler et al., 2017; Bagarinao et al., 2019; Miró-Padilla et al., 2019; Finc et al., 2020). These findings are of significant importance because reorganization of functional networks could increase the efficiency of learning and memory and could even increase the resilience of cognitive processes to damage or deterioration. However, investigating the influence of prior learning on altered functional connectivity in humans is complicated by the diverse cognitive, genetic and lifestyle differences in different individuals.

In the present study, to further elucidate the impact of prior learning on memory related functional connectivity, we have developed a mouse model in which mice are trained in a repeated acquisition spatial learning and memory task for several months. Using mice, we are able to control for environmental and genetic factors, and we can also control for prior learning experiences. Our aim was to investigate whether learning one task would increase the efficiency of the functional networks underlying a different form of memory (contextual fear memory). To do so we adopted a brain-wide functional connectivity approach using immediate early gene imaging that has been recently described (Wheeler et al., 2013; Vetere et al., 2017; Scott et al., 2020).

In this study, we show that chronic cognitive stimulation, in the form of spatial learning is sufficient to induce generalized changes in the organization of functional connectivity networks underlying a test of contextual fear memory.

## Methods

### Mice

8-week-old male C57BL/6J mice purchased from The Jackson Laboratory (Bar Harbour, ME, United States) were used for all experiments. Upon arrival, mice were group housed, 3-4 mice per cage, under a 12-h light/12-h dark cycle with *ad libitum* access to food and water. Behavioural tests and network analyses were conducted using groups of *n* = 10. All procedures were conducted in accordance with protocols approved by the University of Calgary, Health Sciences Animal Care Committee, following the guidelines of the Canadian Council for Animal Care.

### Morris Water Task Training

In order to provide mice with chronic cognitive stimulation, we trained half of the mice on a repeated acquisition and performance testing variant of the Morris Water Task (MWT) (Spanswick et al., 2007). In this version of the task, the hidden escape platform is moved every second day which required the mice to repeatedly acquire new spatial memories throughout a 10-week training period. We used 10 different platform locations, with each platform location occurring on 2 separate occasions. Mice were trained 4 days per week (2 platform locations). During each daily session, mice were given four trials. Each trial lasted a maximum of 60 seconds and was initiated by placing the mouse gently into the pool, facing the wall. The start location was from a different cardinal compass position around the pool for each trial and the order of start locations was randomized each day. Trials were terminated once the mouse located the hidden platform. If the platform was not found after 60 seconds, mice were gently guided to the platform by the experimenter. Once on the platform, mice were given 15 seconds to remain on the platform before being returned to their cage. Trials were interleaved, whereby each mouse performed their first trial before the first mouse performed its second of four daily trials. This resulted in an intertrial interval of approximately 10 minutes. The circular pool had a diameter of 150 cm and a depth of 50 cm. The pool was filled so that the water level was 2 cm above the surface of a circular escape platform that had a diameter of 11 cm. The water was made opaque using white non-toxic tempera paint. The water was kept at a constant temperature of 22°C and stirred and cleaned of debris before each trial. Automated tracking software (ANY-Maze, Stoelting, Wood Dale, IL, United States) was used to record and analyze swim behaviours in the pool, including swim speed, distance travelled and latency to locate the hidden platform.

### Contextual Fear Conditioning

After the conclusion of the Morris Water Task training protocol, all mice were trained in contextual fear conditioning. Training was conducted in sound-attenuated chambers with grated floors through which shocks (0.5 mA; 2 s) were delivered (Ugo Basile, Gemonio, Italy). Mice were first allowed to acclimate to the chamber for 2 minutes prior to the presentation of a series of 3 shocks, each separated by an interval of 90 seconds. 24 h after the training session, mice were returned to the conditioning chambers for a 5-min retention test. During this test, no shocks were administered, and behaviour was monitored via an overhead infrared camera in conjunction with an\ automated tracking software (ANY-Maze, Stoelting, Wood Dale, IL, United States). The chamber was cleaned using 70% ethanol and allowed to dry before and after each trial.

### Perfusions and Histology

Mice were transcardially perfused with 0.1 M phosphate buffered saline (PBS) followed by 4% formaldehyde 90 minutes after retention testing. Brains were then extracted and postfixed in 4% formaldehyde for 24 h. Fixed brains were cryoprotected in 30% W/V sucrose solution at 4°C until no longer buoyant. From cryoprotected brains, serial coronal sections with a thickness of 40 µm were cut on a cryostat (Leica Biosystems, Concord, ON, Canada) and stored in 12 series at -20°C in antifreeze solution.

### Immunohistochemistry

Tissue sections were washed 3 times (10 minutes per wash) in 0.1M PBS before being incubated in a primary antibody solution of 1:2000 rabbit anti-c-Fos primary antibody (226 003, Synaptic Systems, Göttingen, Germany), 3% normal donkey serum, and 0.03% Triton-X100 for 48 h at room temperature on a tissue shaker. Tissue sections were washed 3 × 10 minutes in 0.1M PBS before secondary antibody incubation. The secondary antibody solution was composed of 1:500 donkey anti-rabbit Alexa Fluor 488 (111-545-003, Cedar Lane Labs, Burlington, ON, Canada) in PBS for 24 h at room temperature. Sections were then transferred to 1:2000 DAPI solution for 15 min before being washed 3 × 10 minutes in 0.1M PBS. Labeled sections were mounted to glass slides and coverslipped with PVA-DABCO mounting medium.

### Brain-Wide c-Fos Quantification

Quantification of fluorescent c-Fos labeled cells was conducted using a custom semi-automated segmentation and registration pipeline (Figure 2A). Slides were first imaged using an Olympus VS120-L100-W slide scanning microscope (Richmond Hill, ON, Canada). Images were collected using a 10x objective with a numerical aperture of 0.40 and a Hamamatsu ORCA-Flash4.0 camera. Labelled c-Fos was imaged using a FITC filter cube and a 9.00 V lamp at an intensity of 100% and an exposure time of 140 ms. DAPI staining was imaged under the same conditions, but with a DAPI filter cube and an exposure time of 65 ms. Cells expressing a c-Fos label were segmented using the machine learning-based pixel and object classification program, *Ilastik* (Berg et al., 2019). To further prepare *Ilastik* output images and DAPI channel photomicrographs for regional registration, a custom plug-in was written for *ImageJ*. The pixel intensity threshold of the *Ilastik* outputs was adjusted so as to only contain objects which the program determined to be within the correct range of pixel intensities and shapes. To compensate for inadequate regional area measurements at an image-by-image level in the atlas registration software, a mask of evenly spaced binary points was generated from the DAPI channel image. The pixel intensity thresholds of these images were adjusted to create a binary mask in the shape of the tissue section. Grid lines were then overlaid to create a mask of binary points arranged in a square grid in the shape of the tissue section. Adjacent binary points were spaced by 22 µm, therefore, each point in the mask accounted for an area of 484 µm^2^.

Next, tissue sections were registered to plates of the *Allen Mouse Brain Atlas* using the *R*-based *Whole Brain* software (Fürth et al., 2018). Using this software, DAPI channel images were used as references to which the atlas plates were aligned. The number of segmented c-Fos labeled cells per neuroanatomical region was quantified in Whole Brain. Similarly, the binary point masks were processed to count the number of points in each region. Regional areas were then approximated using a Cavalieri-based point counting approach, whereby the number of mask points in each region was multiplied by the area accounted for by each point. This allowed for the c-Fos labeled cells to be normalized by area and presented as regional cell densities.

### Validation of c-Fos Quantification and Regional Area Approximation

A separate cohort of mice was used for the validation of c-Fos labeled cell segmentation and regional area approximation. c-Fos immunostaining and imaging was identical to the methods described previously. To generate ground truth cell counts as a gold standard for our automated counting procedure, ten 500 µm x 500 µm regions of interest (ROI) were randomly generated from each of several regions including the basolateral amygdala, CA1, dentate gyrus, paraventricular nucleus, and the retrosplenial cortex. These ROIs were processed through our *Ilastik* label segmentation pipeline (representative raw and processed images in Figure 2B). In addition, the same ROIs were hand counted independently by 4 experimenters blind to the automated cell count results. The total numbers of cells counted across all counting boxes were then compared to assess whether or not *Ilastik* could segment fluorescent c-Fos labels within natural and acceptable inter-rater variability.

To assess the accuracy of regional area approximations, areas generated using the Cavalieri-based point counting approach were compared to the areas of these same regions which were manually traced in *ImageJ*. During this analysis, five photomicrographs of each of the following regions were examined: basolateral amygdala, CA1, dentate gyrus, paraventricular nucleus, and the retrosplenial cortex.

### Functional Connectivity Network Generation

We focused on a selection of 97 regions based on our ability to discriminate these regions using a DAPI stained image as reference (see Supplemental Table 1 for list of regions and abbreviations). From this list of regions, c-Fos label densities were cross correlated within each group to generate pairwise correlation matrices. Correlations were filtered by statistical significance and a false discovery rate of 95%. For network analyses, correlation matrices were binarized to adjacency matrices based on Pearson’s correlation coefficient and statistical significance (*r* > 0.9; α = 0.005). This threshold allowed for sufficient network density to study global brain dynamics, while still limiting the analyses to only the strongest and most biologically plausible connections(Schneidman et al., 2006). To ensure that the thresholding parameters did not bias network analyses, additional adjacency matrices were generated using either more (*r* > 0.95; α = 0.0005) or less (*r* > 0.8; α = 0.05) conservative thresholds. To analyze adjacency matrices as network graphs, the 97 neuroanatomical regions were plotted as nodes. Connections were drawn between nodes whereby correlations surpassed correlation matrix thresholding parameters.

### Functional Connectivity Network Analysis

Graph theoretical analyses were applied to network graphs to examine global and local properties of the network. These analyses were guided by the use of the *Brain Connectivity Toolkit* (Rubinov and Sporns, 2010), the *SBEToolbox*(Konganti et al., 2013), and other custom analyses. Network properties examined include *node degree, network density, global efficiency, betweenness centrality, Katz centrality*, and *network resiliency*. In the following definitions, *N* is the array of nodes in the network represented by adjacency matrix *A*. The number of nodes in the network is represented by *n* and the number of connections between nodes is *l*. The variable *a*_*ij*_ is the index into the adjacency matrix which indicates the connection status of nodes *i* and *j*. The presence of a connection is represented by *a*_*ij*_ ≠ 0 (Rubinov and Sporns, 2010).

*Node degree* is the number of connections that link a node to the rest of the network (Rubinov and Sporns, 2010).

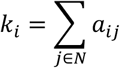

*Network density* is a metric of network dispersion. It is expressed as a proportion of the number of connections in a given network over the number of connections which would be required to saturate a network of the same size (Rubinov and Sporns, 2010).

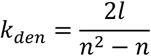

*Global efficiency* is defined as the inverse of the average shortest path of connections between all possible pairs of nodes (Latora and Marchiori, 2001; Achard and Bullmore, 2007; Rubinov and Sporns, 2010).

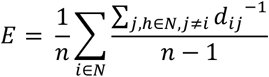

*Betweenness centrality* and *Katz centrality* are measures which can be used to assess the importance of a node in the effective communication of a network. *Betweenness centrality* quantifies the number of shortest paths between nodes that pass-through a given node (Freeman, 1978; Brandes, 2001; Rubinov and Sporns, 2010).

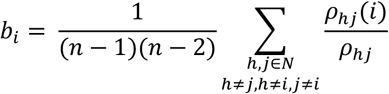

*Katz centrality* applies an eigenvector approach to this metric by weighting the connections involving more highly connected nodes more heavily than those from lesser connected nodes when considering the makeup of the shortest paths which pass through a given node (Katz, 1953; Hubbell, 1965). The attenuation factor, α, used for this analysis was 0.1 (Zhan et al., 2017).

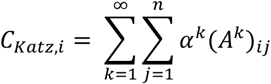

*Network resiliency* was assessed through targeted node deletion and an assessment of the size of the largest community of connected nodes and global network efficiency with each deletion. Nodes were targeted for deletion in decreasing order, from nodes with the highest degree to those with the lowest. Degree was recalculated after each deletion and the list was reordered accordingly.

Network metrics were both compared across conditions as well as used to assess small world-like network properties compared to random control network topology. Small world network distribution can be described as being efficient at both local and global scales (Watts and Strogatz, 1998). Random null control networks were generated for both the cognitive training and control groups and were matched for network size, overall degree, and degree distribution. Local efficiency was assessed by comparing mean clustering coefficients, while global efficiency was assessed by comparing bootstrapped global efficiency values with one hundred replacements.

Networks were considered to display small world-like properties if they had displayed both the high global efficiency characteristic of a random network and increased local efficiency relative to random networks (Wheeler et al., 2013).

### Statistical Analyses

All statistical tests for comparing behavioural data in the Morris Water Maze and contextual conditioning task, c-Fos segmentation and regional area approximation validation, and functional connectivity metric networks were conducted using GraphPad Prism (GraphPad Software, San Diego California US). The analysis of functional connectivity networks was conducted using MATLAB. Figures were generated using MATLAB, Cytoscape, and GraphPad Prism.

## Results

### Morris Water Maze Training Alters Memory Performance in Unrelated Tasks

To assess the generalization of improved cognitive performance following long-term spatial learning (Figure 1A-C), mice were trained and tested in a contextual fear conditioning paradigm (Figure 1D). The percentage of time that mice exhibited freezing behaviour was compared between groups and across the training and retention test sessions. Across both sessions, there were no significant differences in freezing behaviour (Figure 1E). However, when the retention was divided into a first half and a second half, mice who had received cognitive training displayed increased freezing behaviour during the first half of the test and then decreased freezing during the second half of the test (Figure 1F). This may be indicative of changes in memory strength and/or behavioural flexibility.

**Fig. 1:**
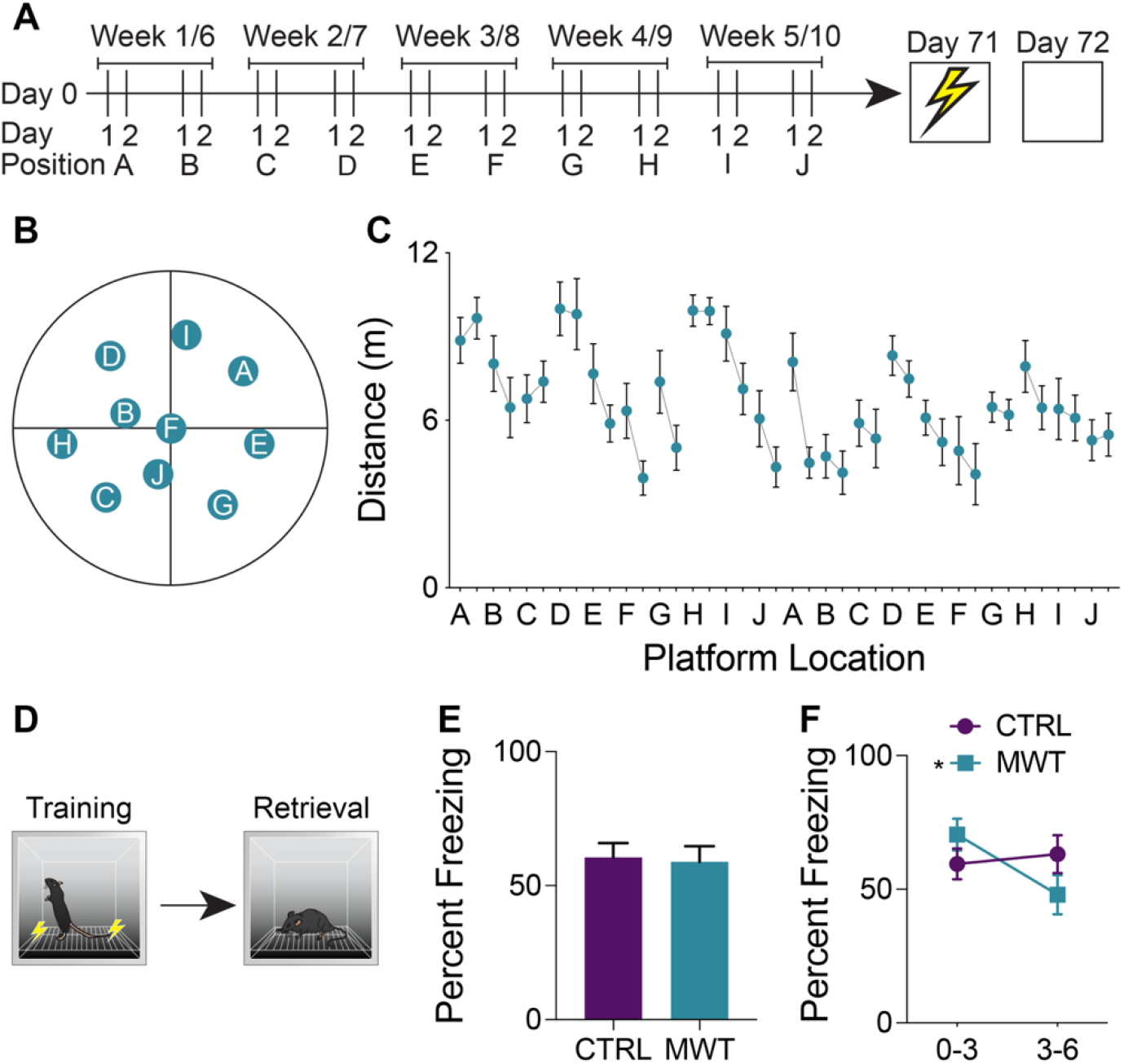
Morris water task training alters memory ability in contextual fear conditioning. (**A**) Mice were trained on a repeated acquisition and performance testing variation of the MWT (*n*=10) or kept in conventional housing conditions (*n*=10) for 10 weeks. Afterwards all mice underwent contextual fear conditioning and a retention test 24 hours later. (**B**) During water maze training, the escape platform was moved every second day between 10 locations. After the 10^th^ position, the platform was returned to the 1^st^ position and the cycle was restarted. (**C**) Mean distance travelled in the water maze across each of the 20 locations. (**D**) During contextual fear memory retrieval, (**E**) there was no overall difference in freezing rates between the control group and the group which had underwent Morris Water Task training (Paired two-tailed t test; *P*>0.05). (**f**F However, mice who had underwent Morris Water Task training froze significantly more during the first half (minutes 0-3) of the test and less during the second half of the test (minutes 3-6; Two-Way Repeated Measures ANOVA; Time × Treatment interaction: F_1,18_=7.763, *p*=0.0122; MWT 0-3 – 3-6: *p*<0.0066). Data shown are mean ± SEM when applicable.

### Validation of c-Fos Segmentation and Neuroanatomical Atlas Registration

To assess the reliability of the semi-automated c-Fos segmentation and mouse brain atlas registration pipeline used in this study (Figure 2A), we compared c-Fos counts obtained using this pipeline to those gathered manually (see representative images Figure 2B). We found that the number of c-Fos labeled cells quantified using *Ilastik* processing fell within the range of values counted manually by four different experimenters across a subset of regions with varying levels of background autofluorescence (Figure 2C). The inter-rater reliability was determined to be 15% across the datasets as a whole and the semi-automated cell counts were within 3.75% of the average of the manual cell counts. Manual and automated cell counts were highly correlated across the sampled brain regions (Figure 2D). *WholeBrain* region registration also produced regional area approximations that correlated highly with hand-traced regional area values (Figure 2E).

**Fig. 2:**
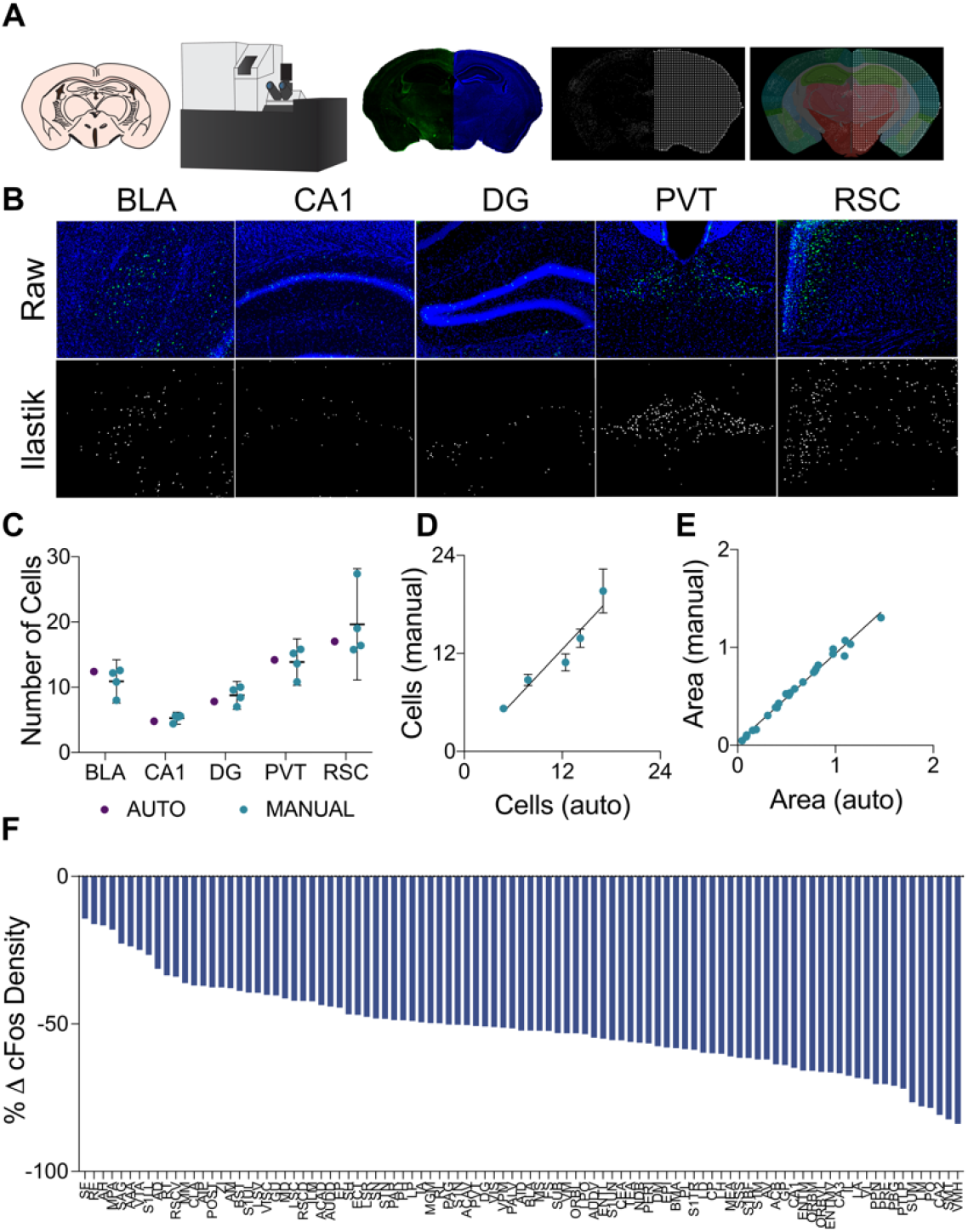
Semi-automated pipeline for generating accurate regional c-Fos densities. (**A**) In the cell segmentation and tissue registration pipeline, tissue was sectioned, immuno-labeled, and mounted on slides prior to being imaged on a fluorescent slide scanning microscope. Images of labelled c-Fos and DAPI staining were then processed using *Ilastik* and *ImageJ* to generate binary c-Fos labels and a mask of evenly spaced grid points in the shape of the tissue sections. These binary images were then applied to plates from the *Allen Mouse Brain Atlas* which had been morphed to align with the tissue sections using *Whole Brain*, yielding regional c-Fos densities. (**B**) Examples of raw c-Fos^+^ cells in the BLA, CA1, DG, PVT, and RSC (top, L-R) and cells segmented using *Ilastik* (bottom). (**C**) A set of ROIs was quantified for validation. Automated *Ilastik* segmentation yielded cell counts within the 95% confidence intervals of counts acquired from trained independent experimenters in each of the aforementioned regions (Two-Way ANOVA; segmentation method factor: F_1,15_=0.09515, *p*=0.7620). Data presented as mean 95% ± confidence interval. (**D**) Cell counts collected using automated *Ilastik* processing were found to correlate highly with mean counts gathered through manual counting (*Pearson r*=0.9610, *p*<0.01). Data presented as mean 95% ± confidence interval. (**E**) Area approximations generated using the pipeline were correlated with areas acquired by tracing regions manually in *ImageJ* (*Pearson r*=0.9941, *p*=<0.0001). (**F**) c-Fos quantification across the 97 brain regions of interest. Relative to the control group, c-Fos expression in the mice that had previously received Morris water task training was decreased in most brain regions.

### Prior spatial learning alters c-Fos expression associated with context memory recall

We quantified regional c-Fos expression density in 97 regions of interest (see Supplementary Table 1 for list of regions). This is depicted in Figure 2F as the percent change from the control condition with long-term spatial learning. Interestingly, we observed a decrease in c-Fos expression density in all brain regions in mice that had received prior spatial training. A relatively small subset of regions showed an increase in c-Fos expression compared to control mice. With respect to brain-wide activity, prior spatial training resulted in an overall significant decrease (Unpaired t test; *p*=0.0003) in c-Fos expression (Supplemental Figure S1).

### Morris Water Maze Training Alters Functional Connectivity Network Topology

Analyses of cross-correlated regional c-Fos expression density revealed differences in global functional connectivity network topology. On a global scale, we observed a reorganization of connections throughout the brain (Figure 3A-F). Relative to control conditions, mice that underwent prior cognitive training exhibited an increase in the overall density of functional connections during subsequent contextual fear memory retrieval (Figure 3G). Furthermore, the organization of these networks after prior cognitive training resulted in increased global efficiency relative to the control condition (Figure 3H). These increases occurred were also present in the networks constructed with both more or less conservative thresholds, indicating that this effect was not an artifact of the thresholding level (Supplemental Figure S2).

**Fig. 3:**
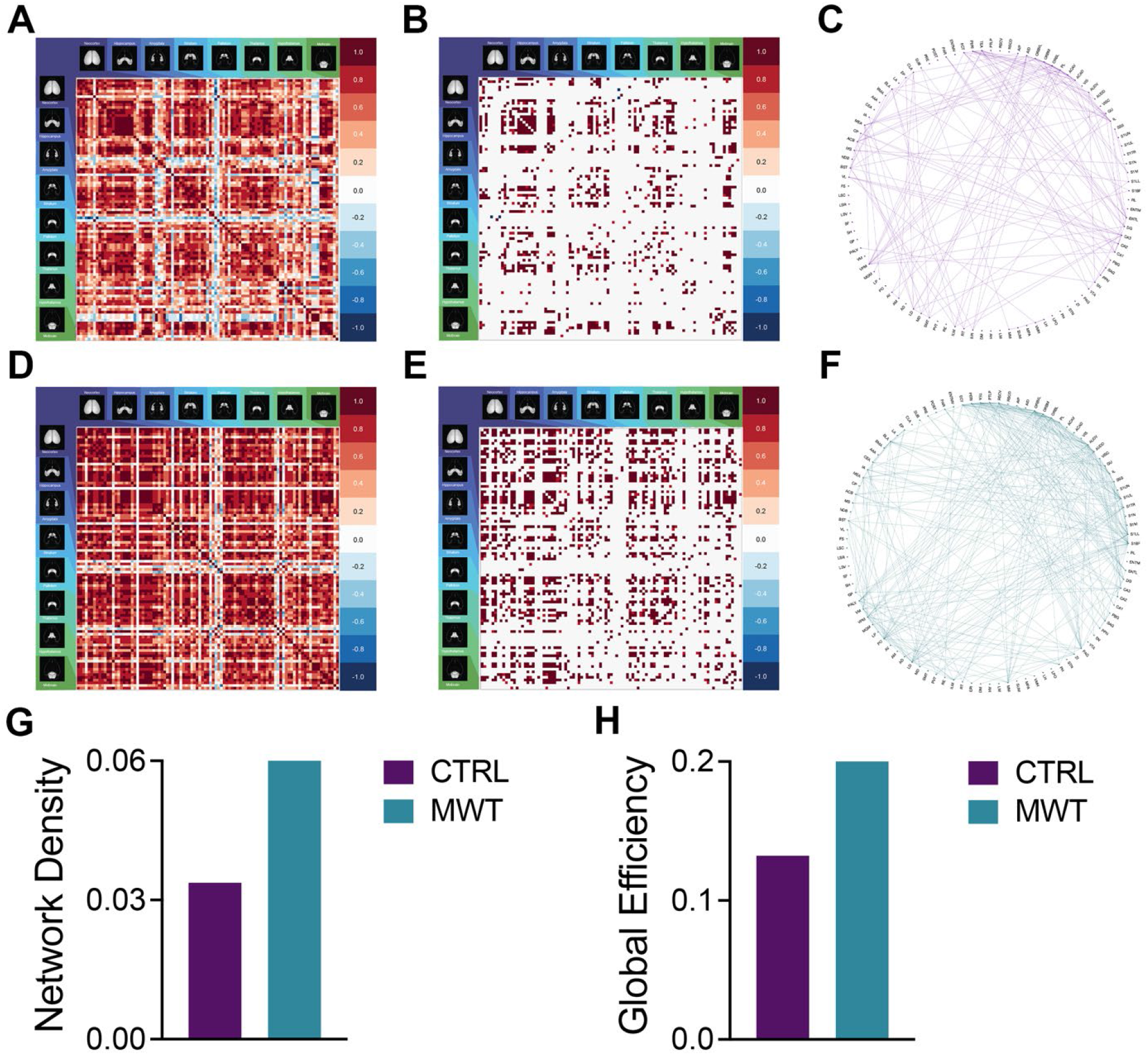
Morris water task training alters global memory network topology. Pairwise correlation matrices for and binarized adjacency matrices and circle plots showing significant correlations between regions for control (**A**-**C**) and Morris water task trained (**D**-**F**) groups. See Online Resource 1 for full list of regions. MWT training increased (**G**) network density and (**H**) global network efficiency.

### Control and Morris Water Maze Training Networks Exhibit Small-World Qualities

Functional connectivity networks in the brain are considered to be complex networks. Many complex networks exhibit small world organization. Small-world networks balance global efficiency with local clustering by having a small proportion of nodes to contribute disproportionately to the overall connectivity of the network (Watts and Strogatz, 1998; Bassett et al., 2006; Sporns and Honey, 2006). This type of organization facilitates specialized processing in dense, local clusters while maintaining efficient information transfer between clusters. Small world organization can be identified in networks with a heavy-tailed degree distribution and increased local clustering without compromise to global efficiency. Both networks from control and spatial learning groups exhibited this heavy-tailed degree distribution, with the majority of nodes making very few connections and a lesser number of nodes carrying disproportionate importance to the overall connectivity of the network (Figure 4A) (Bassett et al., 2006; Bullmore and Sporns, 2009)

**Fig. 4:**
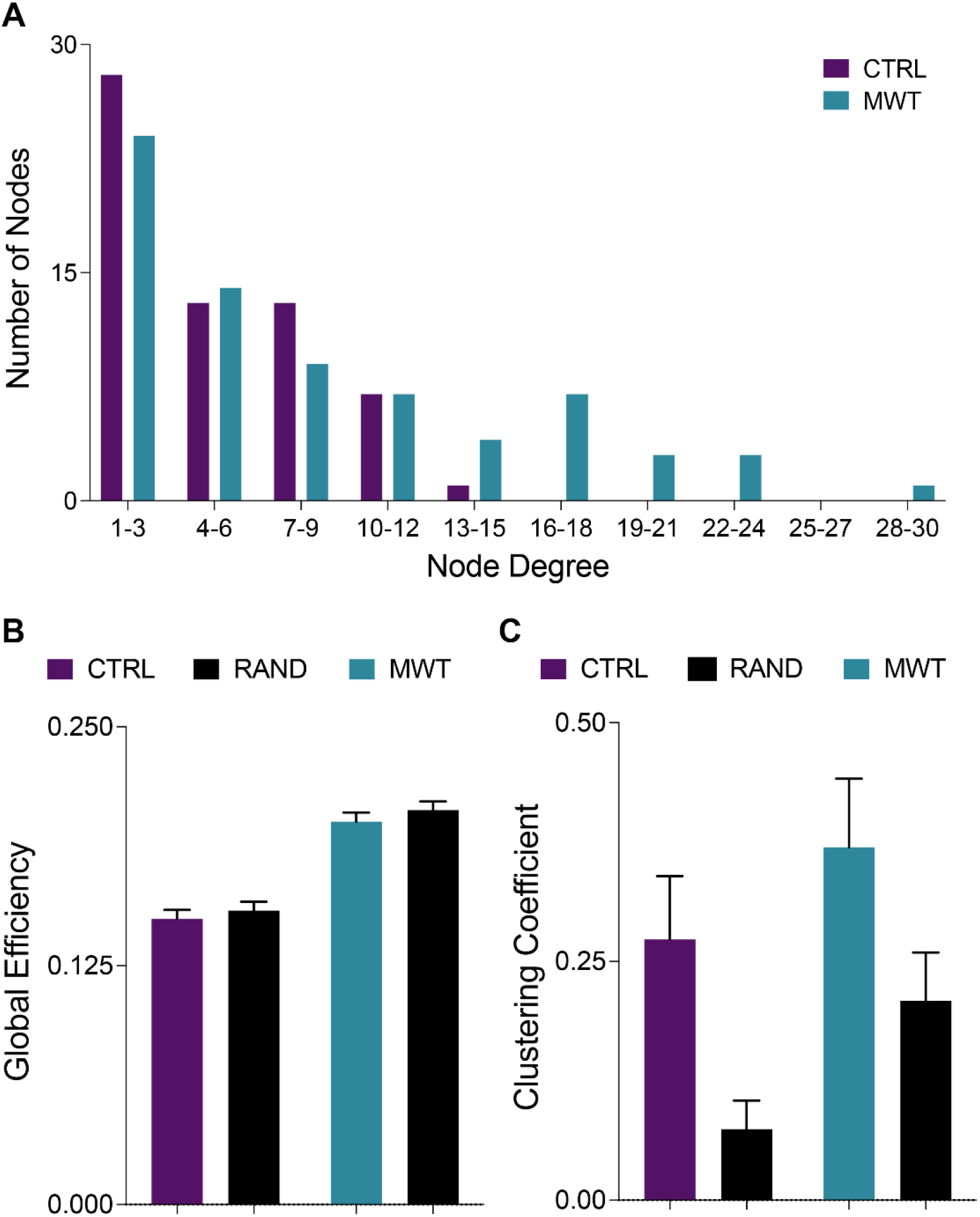
Networks underlying contextual memory recall display small world-like properties. (**A**) Consistent with definitions of small world organization, in both control and MWT trained networks the majority of nodes were of a low degree. However, water maze training shifted the degree distribution and increased the number of highly connected nodes. (**B**) Also consistent with small world organization, both control and MWT trained networks showed equivalent global efficiency to random networks matched for degree distribution. (**C**) A third requirement for the classification of a small world organization is a higher clustering coefficient than a random network. Compared to random networks matched for degree distribution, both control and MWT trained networks showed heightened clustering. Data shown are mean ± 95% confidence intervals.

Additional small-world properties based on clustering and efficiency were assessed by comparing functional connectivity networks to randomly generated control networks. These randomly generated networks were matched for size and degree distribution (Rubinov and Sporns, 2010). The random networks demonstrate the extent to which clustering would be expected in a network based purely on chance alone (Sporns, 2018). This then indicates that differences in topological characteristics between observed biological networks and the random null models are core to the behaviour of the biological networks (Betzel et al., 2016). Interpretations of comparisons to randomly generated null networks must consider that random networks have low mean clustering, which is associated with low local efficiency. However, while random network organization yields a low mean clustering coefficient, they also are characterized by having short mean path lengths, which is reflected in a high global efficiency (Watts and Strogatz, 1998).

Comparisons to random null networks highlighted that both the control and spatial learning networks showed increased clustering (Figure 4B) while maintaining the high global efficiency characteristic of random networks (Figure 4C). Together, these analyses indicate that the functional connectivity networks engaged during memory recall in both the control and spatial learning conditions exhibit properties that are consistent with small-world topology.

### Morris Water Maze Training Alters Cluster Organization and Connectivity

Changes in network topology were also observed at the local level. The organization of local communities within global networks changes with long-term spatial learning. While the size of the giant component (GC) (Figure 5A-C) underwent very little change with MWT training, differences arose in the connectivity patterns within this component. Within the GC, MWT training increases the mean number of connections per node (Figure 5D). The changes in connectivity coincided with changes in network resiliency. When faced targeted deletion of nodes, with deletions occurring in the order of decreasing degree, long-term spatial learning increased the ability of the network to preserve its global efficiency (Figure 5E) and giant component size (Figure 5F).

**Fig. 5:**
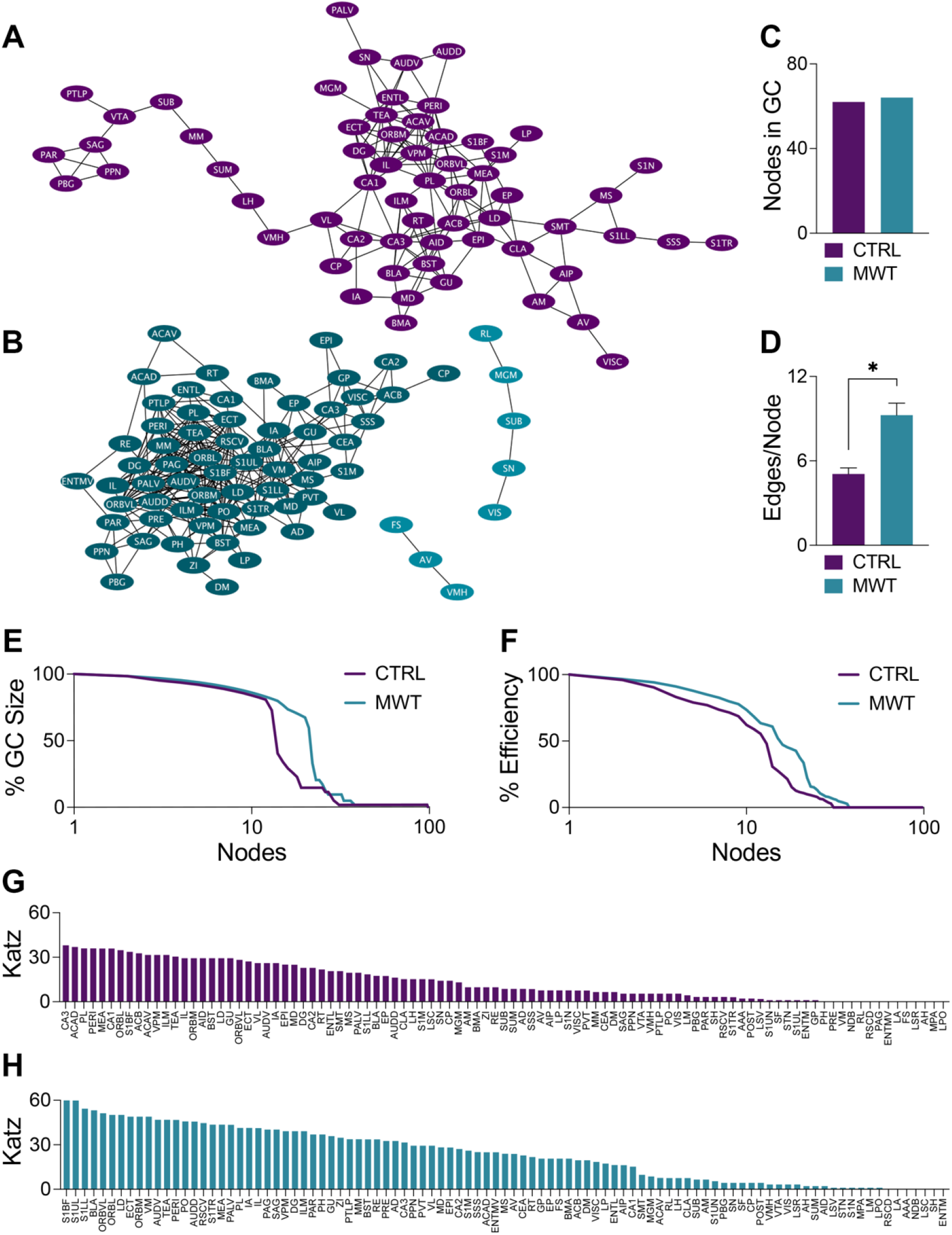
Morris water task training alters clustering patterns and increases network resiliency. Network plots of (**A**) control and (**B**) MWT trained networks with the giant component (GC) highlighted in each in the darker shade. (**C**) In these networks, there is very little difference in the size of the GC. (**D**) Within the GC, there was an increase in the mean number of edges per node with MWT training (Two-tailed t test, *p*<0.0001). MWT training also made the (**E**) global efficiency and (**F**) integrity of the GC more resilient to targeted node deletion. Relative to the control condition (**G**), MWT training (**H**) also increased Katz centrality of a subset of regions within the network. Data shown are mean ± SEM when applicable.

Coinciding with these changes in network resiliency were changes in Katz centrality. Katz centrality is a measure of centrality which differentially weighs connections based on the degrees of the nodes involved (Katz, 1953). This measure has previously been shown to correlate highly with neuronal activity compared to other measures of centrality (Fletcher and Wennekers, 2018). While most nodes in the control network had similar Katz centrality vectors (Figure 5G), MWT training increased the centrality of a subset of regions (Figure 5H). There was considerable overlap between these regions with increased Katz centrality and the regions in the most densely connected region of the GC. This was further corroborated by analysis of regional degree distribution (Supplemental Figure S3) and change in Katz centrality (Supplemental Figure S4) which further highlighted an increase in connectivity and of importance of numerous amygdala subregions following spatial learning.

## Discussion

In the current study we employed a brain-wide activity mapping approach to examine the impact of a prolonged period of repeated spatial learning on brain-wide patterns of activation and functional connectivity. We posited that repeated activation of the circuits underlying spatial learning and memory might alter the networks that represent other forms of hippocampus dependent memory in the future. Our results clearly indicate that this prior spatial learning manipulation caused significant changes to the task-related functional connectivity associated with retrieval of a contextual memory. In the current study, prior spatial training had minimal effects on the overall retrieval of a subsequently acquired contextual fear memory. Interestingly, when subdividing the retrieval period, we noticed that the nature of the memory was different in mice that had prior spatial training. Specifically, the retention was stronger in the first half of the test and decreased in the second half relative to control mice. This could be indicative of an increase in behavioural flexibility and/or increased rate of extinction in the absence of additional foot shocks. That we did not observe major differences in memory retrieval was not surprising given that the subjects were normal mice without memory deficits, the memory in control mice was already very strong and the retention interval was short (24 hours). This similarity in behavioural performance between conditions allowed us to assess patterns of neuronal activation without the confound of differential memory ability. When we examined neuronal activation and network organization underlying the memory in these two groups, we observed a number of differences that could enhance retention/retrieval in the face of cognitive decline.

We were most interested in investigating whether such a manipulation, which might be viewed as memory practice or training, would enhance measures of efficiency when examining the storage and retrieval of future memories. The efficiency of brain activity underlying cognitive function is vulnerable to aging and disease. Decreased efficiency of cognitive processing has been reported in several neurological conditions, including major depressive disorder (Zhang et al., 2020), schizophrenia (Sheffield et al., 2016), and Alzheimer’s disease (Srivishagan et al., 2020). Even in healthy adults, patterns of brain activation become less efficient with age (Ajilore et al., 2014; Chong et al., 2019). These decreases in efficiency also coincide with decreased cognitive performance, thereby indicating that an intervention which can improve the efficiency of the functional connectome may preserve cognitive function in these conditions (van den Heuvel et al., 2009). We show here that the efficiency of brain-wide activation, as measured by c-Fos expression, is greatly increased in the mice that had prior spatial training. Efficiency can be defined as equal or greater memory performance with the expenditure of fewer resources (i.e., a decrease in activation) (McQuail et al., 2020). Our results show that contextual memory retrieval following spatial learning was associated with a decrease in the total c-Fos expressing cells throughout the brain compared to mice that had not previously experienced any spatial training. When examining this activation regionally we observed that all brain regions exhibited reduced activity. This pattern of activity perhaps indicates that the behavioural expression of the memory retrieval is more efficient on the level of neuronal activation.

At a network level, global efficiency can be estimated as the inverse of the average shortest path lengths between all network nodes. This measure represents the relative ease or difficulty of integrating information between nodes in a network. Using this measure, we found that mice which had received prior spatial training exhibited enhanced global efficiency and higher clustering compared to controls during subsequent contextual memory retrieval. Both of these findings are consistent with the effects of memory training observed using human functional neuroimaging(Langer et al., 2013). This analysis corroborates the interpretations of increased efficiency based on overall brain activation. Further corroborating these interpretations are changes in the Katz centrality within these networks. Centrality measures can be used as proxies of the relative importance of a node in the maintenance of effective communication across a network and as indicators of the functional segregation of the network as a whole. Spatial training increased the centrality of a subset of nodes. This pattern of distribution suggests a higher degree of network segregation and that this subset of regions is relatively more important to the behavioural expression of the context memory. Comparatively, based on the variability of regional Katz centrality, all regions in the network generated from untrained mice are of similar importance in the expression of this same behaviour. These findings coincide with increased centrality in resting state memory networks following working memory training in human neuroimaging studies(Takeuchi et al., 2017). Together these metrics illustrate that the redistribution of neuronal activation induced by prior spatial learning is not only more efficient from the perspective of energetic resources, but also proves to be more efficient with respect to global flow of information throughout the brain.

Regardless of prior spatial training exposure, the networks engaged by contextual memory retrieval displayed characteristic small-world properties. Compared to random networks, we observed that the memory networks had a small number of highly connected regions, an increase in clustering coefficient, and equivalent global efficiency. Small world organization facilitates specialized processing in densely connected local communities while also allowing for efficient transfer of information between local communities. We used Markov chain clustering to detect the structure of these communities within the networks. In both networks there was a large interconnected central component, referred to as the giant component, which did not change in size as a result of prior spatial training. However, there was a significant increase in the density of connections within the GC of the group of mice who had received prior spatial learning. The densely connected giant component at the core of the network underlying context memory expression in mice with prior spatial training contained many redundant connections. Therefore, we hypothesized that more successive deletions would be required to break apart communities in a way which would be consequential to the effective communication of the network. This hypothesis was supported by the results of targeted node deletion. By sequentially deleting nodes in the descending order of their degree, we noted that mice which had prior spatial training were able to retain a higher percentage of their basal global efficiency and giant component size. In being more resistant to targeted attack, this network can be said to be more resilient than the network obtained from control mice (Achard et al., 2006).

Increased resiliency to targeted node deletion presents interesting possibilities from the perspective of neurodegenerative disease. The pathology of many neurodegenerative conditions does not arise uniformly throughout the brain and rather targets the most highly involved regions of a network (reviewed in (Crossley et al., 2014). In addition to suffering from targeted attacks, the functional connectomes characteristic of many neurodegenerative conditions networks also display decreased redundancy in their connectivity patterns, rendering networks more vulnerable to these attack (Langella et al., 2021). An efficient network which is more resilient to attack has the potential to delay or reduce cognitive decline during early neurodegenerative disease progression (Rittman et al., 2019). The present study found that cognitive stimulation through repetitive learning experiences was able to increase network efficiency and resilience. Therefore, the potential exists for prior exposure to repetitive learning experiences to increase resiliency to deterioration. Future studies of this phenomenon might build upon this by examining whether prolonged cognitive stimulation and the resulting alterations in functional connectivity are sufficient for reducing cognitive deficits observed in early stages of neurodegeneration.

In the present study, networks were generated based on correlated expression of c-Fos across the brain. While this method has been demonstrated at various levels of regional organization in previous publications(Wheeler et al., 2013; Vetere et al., 2017; Silva et al., 2019; Scott et al., 2020), it is worth acknowledging the limitations of this approach. While exhibiting excellent spatial resolution at the single cell level, c-Fos expression is limited in its temporal resolution. Consequently, it is impossible under the current design to establish patterns of c-Fos expression during distinct bouts of freezing or movement during conditioned context reintroduction. In an experiment such as this in which freezing behaviours were consistent between groups, it is possible that this limitation has less of an impact on the ability to interpret the results than in an experiment in which the behaviours corresponding with the tagged neuronal activity vary greatly between groups. In such scenarios, follow-up experiments using an *in vivo* measure of regional activation would be advised to assess the specificity functional connections to distinct behavioural outputs.

Additionally, while we attribute the changes in network topology to spatial learning, there are other possible factors, such as stress and exercise, present during these episodes which may also contribute to network reorganization. Under the current design, it was decided that measures controlling for exercise, such as the use of a yoked control group, would fail to control for stress and vice versa. Furthermore, the long-term nature of the learning protocol limited the design to tasks which could provide repeated memory acquisition. Future studies might choose to assess this paradigm using cognitive tasks which are less reliant on animal movement and may also be less stress-inducing.

## Acknowledgements

Funding for this study was provided by The Brain Canada Foundation/The Azrieli Foundation Early Career Capacity Building Grant (4709) and an NSERC Discovery Grant (RGPIN-2018-05135) to JRE. We acknowledge the Hotchkiss Brain Institute Advanced Microscopy Platform and the Cumming School of Medicine for support and use of the Olympus VS120-L100-W slide scanning microscope.

## Declarations

### Availability of Data and Material

Data accompanying the figures in the manuscript has been provided (Supplemental Data1). Additional raw data will be made available upon request.

### Code Availability

Code used during analyses have been uploaded to the following GitHub repository: https://github.com/dterstege/PublicationRepo/tree/main/Terstege2022A

### Authors’ Contributions

DJT and JRE conceived of and designed the experiment. DJT and IMD conducted experiments. DJT and JRE conducted analyses and wrote the manuscript.

### Ethics Approval

All procedures were conducted in accordance with protocols approved by the University of Calgary, Health Sciences Animal Care Committee, following the guidelines of the Canadian Council for Animal Care.

