## Supplemental Figures for "Brain-wide neuronal activation and functional connectivity are modulated by prior exposure to repetitive learning episodes"

Supplemental Information

Dylan J. Terstege[1] Isabella M. Durante[1] & Jonathan R. Epp[1]*

[1]Department of Cell Biology and Anatomy, Hotchkiss Brain Institute, Cumming School of Medicine, University of Calgary, 3330 Hospital Drive NW, Calgary, Alberta, Canada T2N 4N1.

* Corresponding author
Dept. of Cell Biology & Anatomy
University of Calgary
HMRB 162, Health Sciences Centre
3330 Hospital Dr. NW
Calgary, AB
T2N 4N1


**Supplementary Table 1: List of brain regions (nodes) in which c-Fos was quantified. Numbers and color coding represent the order and subdivisions in which they appear in the correlation matrices.**

| # | Abbreviation | Full Name | # | Abbreviation | Full Name |
|---|---|---|---|---|---|
| 1 | S1bf | Primary somatosensory area, barrel field | 49 | CP | Caudoputamen |
| 2 | S1ll | Primary somatosensory area, lower limb | 50 | ACB | Nucleus accumbens |
| 3 | S1m | Primary somatosensory area, mouth | 51 | FS | Fundus of striatum |
| 4 | S1n | Primary somatosensory area, nose | 52 | LSc | Lateral septal nucleus, caudal part |
| 5 | S1tr | Primary somatosensory area, trunk | 53 | LSr | Lateral septal nucleus, rostral part |
| 6 | S1ul | Primary somatosensory area, upper limb | 54 | LSv | Lateral septal nucleus, ventral part |
| 7 | S1un | Primary somatosensory area, unassigned | 55 | SF | Septofimbrial nucleus |
| 8 | SSs | Supplemental somatosensory area | 56 | SH | Septohippocampal nucleus |
| 9 | IL | Infralimbic area | 57 | GP | Globus pallidus |
| 10 | GU | Gustatory areas | 58 | PALv | Pallidum, ventral region |
| 11 | VISC | Visceral areas | 59 | MS | Medial septal nucleus |
| 12 | AUDd | Dorsal auditory area | 60 | NDB | Diagonal band nucleus |
| 13 | AUDv | Ventral auditory area | 61 | BST | Bed nucleus of the stria terminalis |
| 14 | VIS | Visual areas | 62 | VL | Ventral lateral nucleus of the thalamus |
| 15 | ACAd | Anterior cingulate area, dorsal part | 63 | VM | Ventral medial nucleus of the thalamus |
| 16 | ACAv | Anterior cingulate area, ventral part | 64 | VPM | Ventral posteromedial nucleus of the thalamus |
| 17 | PL | Prelimbic area | 65 | MGM | Medial geniculate complex, medial part |
| 18 | ORBl | Orbital area, lateral part | 66 | LP | Lateral posterior nucleus of the thalamus |
| 19 | ORBm | Orbital area, medial part | 67 | PO | Posterior complex of the thalamus |
| 20 | ORBvl | Orbital area, ventrolateral part | 68 | AV | Anteroventral nucleus of the thalamus |
| 21 | AId | Agranular insular area, dorsal part | 69 | AM | Anteromedial nucleus of the thalamus |
| 22 | AIp | Agranular insular area, posterior part | 70 | AD | Anterodorsal nucleus of the thalamus |
| 23 | RSCd | Retrosplenial area, dorsal part | 71 | LD | Lateral dorsal nucleus of the thalamus |
| 24 | RSCv | Retrosplenial area, ventral part | 72 | MD | Medial dorsal nucleus of the thalamus |
| 25 | PTLp | Posterior parietal association areas | 73 | SMT | Submedial nucleus of the thalamus |
| 26 | TEa | Temporal association areas | 74 | PVT | Paraventricular nucleus of the thalamus |
| 27 | PERI | Perirhinal area | 75 | RE | Nucleus of reuniens |
| 28 | ECT | Ectorhinal area | 76 | ILM | Intralaminar nuclei of the dorsal thalamus |
| 29 | CA1 | Ammon's horn, CA1 field | 77 | RT | Reticular nucleus of the thalamus |
| 30 | CA2 | Ammon's horn, CA2 field | 78 | EPI | Epithalamus |
| 31 | CA3 | Ammon's horn, CA3 field | 79 | DM | Dorsomedial nucleus of the hypothalamus |
| 32 | DG | Dentate gyrus | 80 | AH | Anterior hypothalamic area |
| 33 | ENTl | Entorhinal area, lateral part | 81 | LM | Lateral mammillary nucleus |
| 34 | ENTm | Entorhinal area, medial part | 82 | MM | Medial mammillary nucleus |
| 35 | ENTmv | Entorhinal area, medioventral part | 83 | SUM | Supramammillary nucleus |
| 36 | PAR | Parsubiculum | 84 | MPA | Medial preoptic area |
| 37 | POST | Postsubiculum | 85 | VMH | Ventromedial hypothalamic nucleus |
| 38 | PRE | Presubiculum | 86 | LH | Lateral hypothalamic nucleus |
| 39 | SUB | Subiculum | 87 | LPO | Lateral preoptic nucleus |
| 40 | CLA | Claustrum | 88 | PH | Posterior hypothalamic nucleus |
| 41 | EP | Endopiriform nucleus | 89 | STN | Subthalamic nucleus |
| 42 | LA | Lateral amygdalar nucleus | 90 | ZI | Zona incerta |
| 43 | BLA | Basolateral amygdalar nucleus | 91 | PAG | Periaqueductal gray |
| 44 | BMA | Basomedial amygdalar nucleus | 92 | VTA | Ventral tegmental area |
| 45 | AAA | Anterior amygdalar area | 93 | SN | Substantia nigra |
| 46 | CEA | Central amygdalar nucleus | 94 | PPN | Pedunculopontine nucleus |
| 47 | IA | Intercalated amygdalar nucleus | 95 | SAG | Nucleus sagulum |
| 48 | MEA | Medial amygdalar nucleus | 96 | PBG | Parabigeminal nucleus |
| | | | 97 | RL | Rostral linear nucleus raphe |

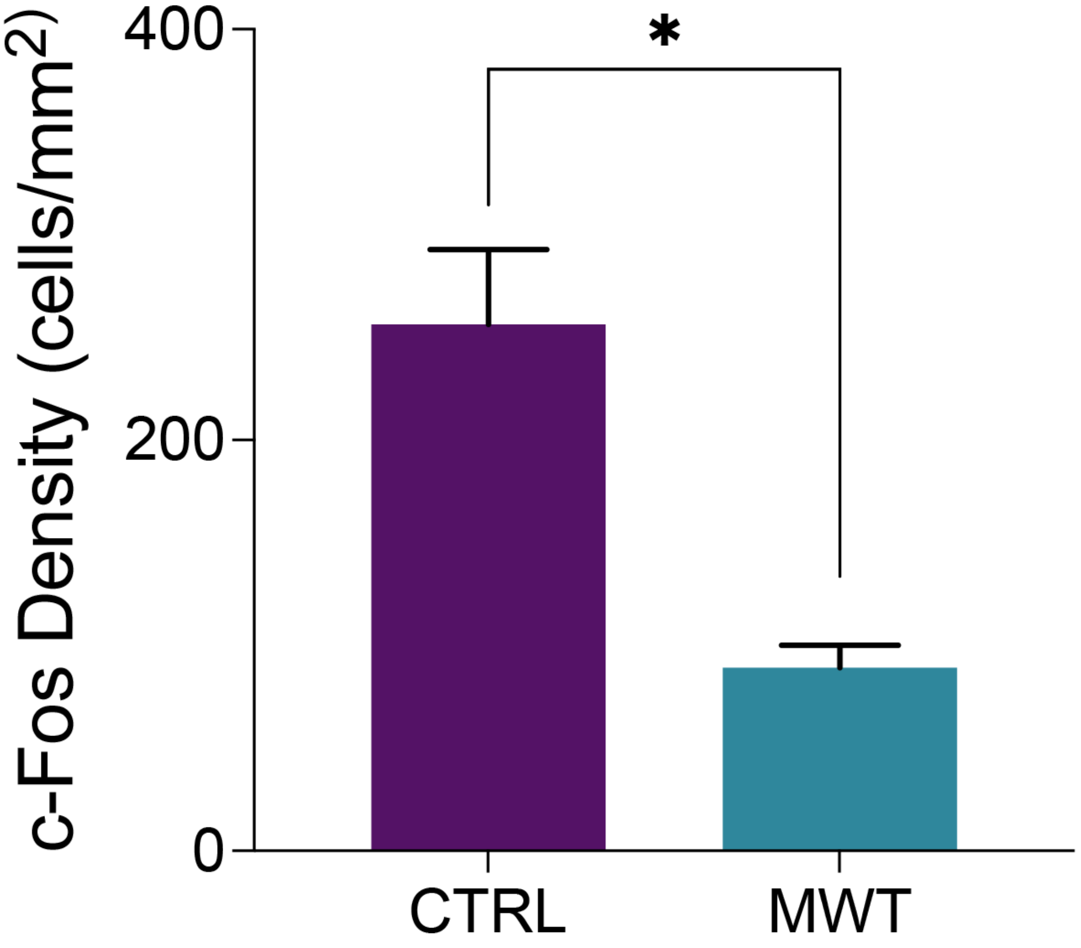


**Supplementary Figure S1: Morris water task training decreased the total brain c-Fos expression density.** The overall c-Fos expression density calculated from across the brain was significantly decreased with MWT training (Two-tailed t test, $p$=0.0003). Data shown are mean ± SEM.

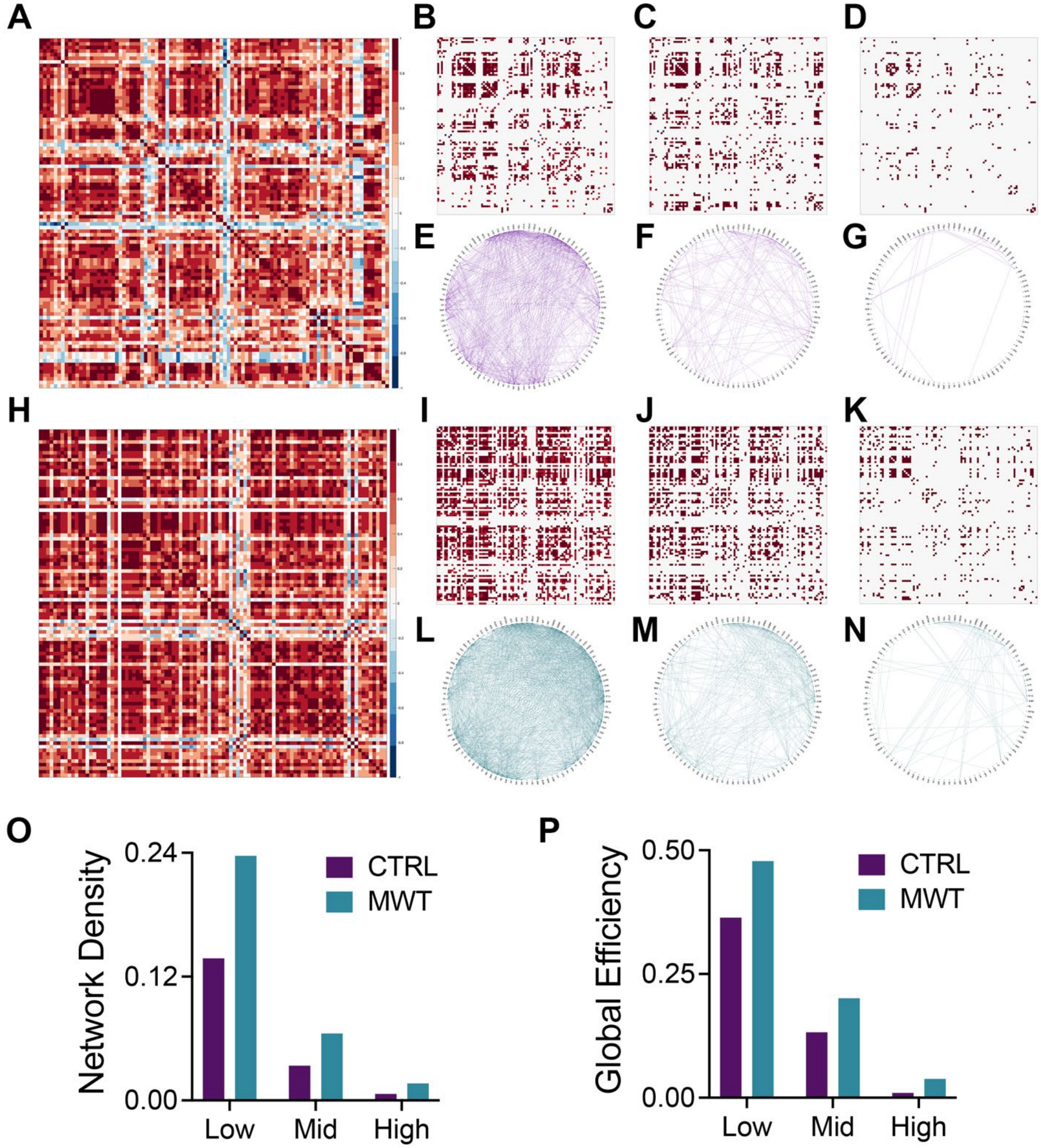


**Supplementary Figure S2: Altered global memory network topology induced by Morris water task training is stable across multiple binary network thresholds.** Pairwise correlation matrices for and binarized adjacency matrices and circle plots showing significant correlations between regions for control (**A**-**G**) and Morris water task trained (**H**-**N**) groups. Networks were binarized at three different confidence thresholds of $R>0.80$, $P<0.05$ (**B**, **E**, **I**, **L**), $R>0.90$, $P<0.005$ (**C**, **F**, **J**, **M**), and $R>0.95$, $P<0.0005$ (**D**, **G**, **K**, **N**). Across all network thresholds, MWT training increased (**O**) network density and (**P**) global network efficiency. See Online Resource 1 for full list of regions.

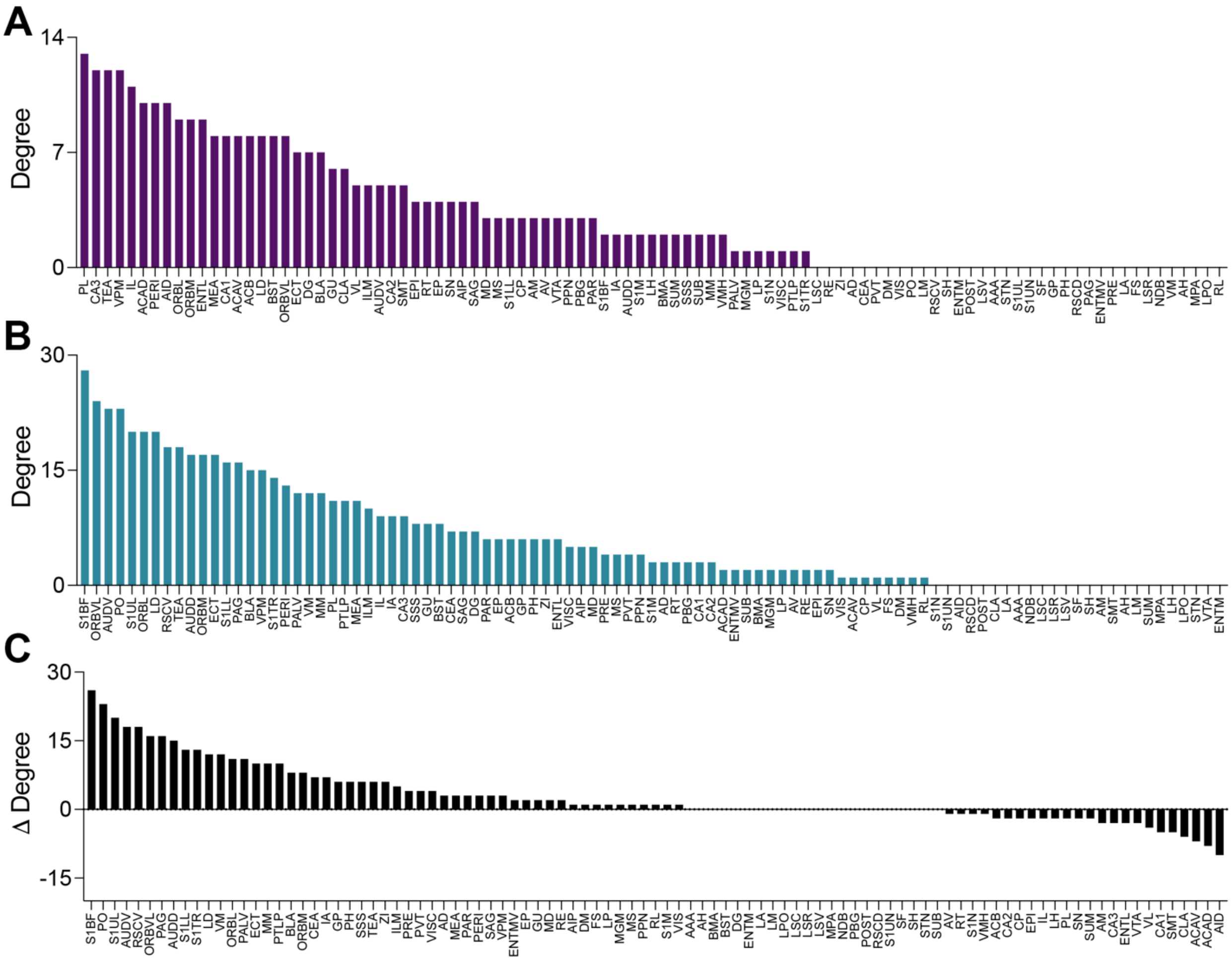


**Supplementary Figure S3: Morris water task training alters degree distribution.**
The degree (i.e., number of functionally connected regions) of each region from both the (**A**) control and (**B**) MWT trained conditions. (**C**) The relative change in the degree of each region in trained mice compared to control mice.

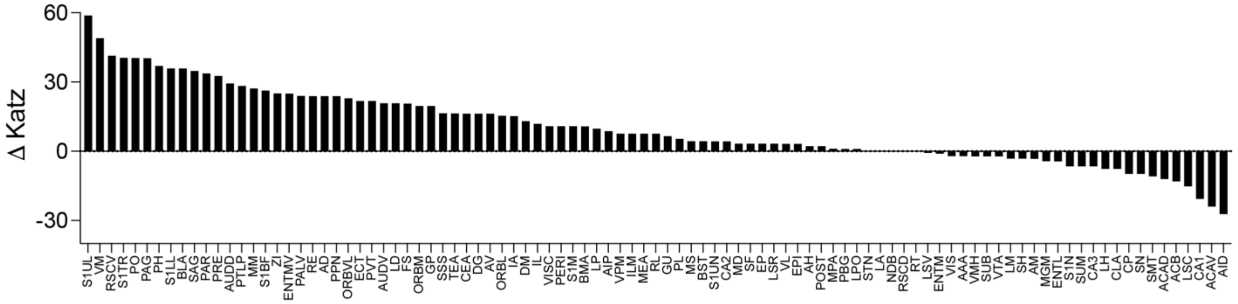


**Supplementary Figure S4: Change in Katz centrality.** The difference in Katz centrality in Morris water task trained mice relative to untrained controls. Most regions are similar between conditions but there is a small number of regions that show a large increase.
